# The co-chaperone DNAJA2 buffers proteasomal degradation of cytosolic proteins with missense mutations

**DOI:** 10.1101/2024.02.15.580437

**Authors:** Heather A Baker, Jonathan P Bernardini, Veronika Csizmók, Angel Madero, Shriya Kamat, Hailey Eng, Jessica Lacoste, Faith Au Yeung, Sophie Comyn, Gaetano Calabrese, Brian Raught, Mikko Taipale, Thibault Mayor

## Abstract

Mutations can result in the loss of a protein’s native function due to protein misfolding, which is generally handled by an intricate protein quality control network. To better understand the triaging mechanisms of misfolded cytosolic proteins, we screened a human mutation library to identify a panel of unstable mutations. The degradation of these mutated cytosolic proteins is largely dependent on the ubiquitin proteasome system. Using BioID proximity labelling, we found that the co-chaperones DNAJA1 and DNAJA2 are key interactors of one of the mutated proteins. Notably, the absence of DNAJA2 increases the turnover of the mutant protein but not of the wild-type protein. Our work indicates that missense mutations in cytosolic proteins can promote interactions with molecular chaperones that normally do not occur. Assessment of the broader panel of cytosolic mutant proteins shows that the co-chaperone DNAJA2 exhibits three distinct behaviours: acting to stabilize solely the mutant, both the wild-type and mutant proteins, or being dispensable. Our work illustrates how distinct elements of the protein homeostasis network are utilized in the presence of a cytosolic misfolded protein.

**Summary Statement:** We identified a panel of cytosolic mutant proteins degraded by the proteasome. DNAJA2 is often required to prevent mutant protein turnover, even if it is sometimes dispensable for the wild-type protein.

## Introduction

Normal cellular function is contingent on the proper folding of newly translated proteins, protein localization, assembly of protein complexes, refolding of misfolded proteins, and degradation of aberrant proteins. Proteostasis, or proteome homeostasis, finely regulates and balances these processes to maintain the proteome and cellular function. The consequences that can arise from the dysfunction of proteostasis have implications for several degenerative diseases and pathological states linked to protein aggregation and misfolding (Yerbury et al., 2016).

Mutations in proteins can induce misfolding, challenge proteostasis, and trigger several protein quality control (QC) mechanisms. Notably, missense mutations are among the most common sequence changes in Mendelian disorders, accounting for more than half of all reported mutations in the Human Gene Mutation database (Sahni et al., 2015). Examples of diseases characterized by missense mutations include cystic fibrosis, in which mutations in the cystic fibrosis transmembrane regulator (CFTR) gene lead to protein misfolding, mislocalization and degradation resulting in a loss-of-function phenotype (Kopito, 1999; Ward et al., 1995). There are thousands of other rare genetic diseases characterized by missense mutations (Stefl et al., 2013). As extensive protein QC mechanisms have evolved to mitigate the potentially harmful effects associated with protein misfolding, mutations causing misfolding can often lead to a loss of function phenotype due to proteolysis.

Protein QC is a highly regulated process that is compartmentalized within eukaryotic cells. Different cellular compartments have specialized proteins that perform distinct functions in maintaining proteostasis and preventing the accumulation of misfolded proteins in the cell (Sontag et al., 2017). A well-characterized example of this is the endoplasmic reticulum-associated degradation (ERAD) pathway that is linked to the ubiquitin proteasome system (UPS). The UPS is one of the main degradation pathways used by the cell whereby an enzymatic cascade of the E1 activating enzymes, E2 conjugating enzymes, and E3 ligases allows for the ATP-dependent tagging of errant proteins with poly-ubiquitin chains and subsequent degradation at the 26S proteasome. HRD1 and AMFR are viewed as the main mammalian E3 ligases responsible for broad substrate recognition of ERAD substrates (Olzmann et al., 2013). In addition, the endoplasmic reticulum has a dedicated chaperone system consisting of BiP, calnexin, calreticulin, and other co-chaperones, that assists in the folding and QC of newly synthesized proteins before they are transported to their final destinations (Needham et al., 2019). In the mammalian cytoplasm, the UPS is also employed in quality control (Baker and Bernardini, 2021). Several E3 ligases have been implicated in the QC of cytosolic proteins, as well as the removal of unassembled components of multiprotein complexes (Chu et al., 2013; Fang et al., 2014, 2011; Haakonsen et al., 2024; Heck et al., 2010; Murata et al., 2001; Nguyen et al., 2017; Xu et al., 2016; Yanagitani et al., 2017; Yau et al., 2017; Zavodszky et al., 2021). Numerous molecular chaperones interact with QC E3 ligases to help mediate proteolysis forming an intricate cellular “circuitry” (Kevei et al., 2017). A significant question that remains is how protein QC mechanisms triage misfolded cytosolic proteins. Especially, the specific identities and roles of chaperones and other quality control proteins that target proteins with missense mutations have yet to be determined.

To better understand the network of QC proteins that are recruited when cytosolic proteins misfold, we identified proteins with missense mutations that are unstable relative to their wild-type (WT) counterparts. One of these model substrates was then used in a BioID proximity labelling experiment to identify which proteins were recruited to the misfolded substrate but not the WT protein. We then showed that DNAJA2, an Hsp40/J-domain protein identified using the BioID approach, is a key QC element that stabilizes some cytosolic misfolded proteins with missense mutations.

## Results

### Identification of unstable variants of cytosolic proteins

To examine protein quality control pathways that target cytosolic misfolded proteins for degradation, we first sought to identify new model substrates by screening a human mutation collection (hmORFeome1.1) for variants with missense mutations that cause proteolysis (Sahni et al., 2015). Notably, by re-analyzing the data from a recent study (Lacoste et al., 2023), we identified over 600 variants expressed at lower levels when expressed in HeLa cells, including 152 cytosolic proteins (Figure 1B). Over 30% of proteins with attenuated expression levels also displayed a significantly increased interaction with elements of the protein QC network (Figure 1C) (Sahni et al., 2015). To confirm the lower signal was due to proteolysis, we selected 11 variants of cytosolic proteins with lower immunofluorescence (IF) intensity, including 7 variants with increased interaction with the protein QC network (Table 1, S2). We also selected another 7 variants that either displayed an increased interaction with elements of the protein QC network and/or a lower stability in a previous study (Sahni et al., 2015). Candidate mutants and their corresponding WT genes were cloned into a bicistronic dual fluorescent reporter vector to assess fluorescence by flow cytometry (Figure 1D)(Yen et al., 2008). In this configuration, the red fluorescent DsRed is used to control the expression level in a given cell, while the signal of the green fluorescent protein EGFP that is expressed downstream of an internal ribosome entry site (IRES) is used to measure the relative level of a variant or its corresponding WT gene tagged C-terminally. Remarkably, all of the 18 assessed mutant reporters display a decrease in green versus red fluorescence in comparison to their WT counterparts (Figure 1E, F), indicating that the variants are likely degraded. 17/18 of these missense mutations are located within a secondary structure, including 12 mutations located within an α-helix (Table 1). Four of the point mutations are to proline residues, while eight of the point mutations result in a change in the charge relative to the WT variant (Table S2). Of the 18 missense mutations, 14 are in a buried region of the protein structure, therefore lower levels of the mutant variant are likely the result of the destabilization of the tertiary structure of these proteins.

**Figure 1.**
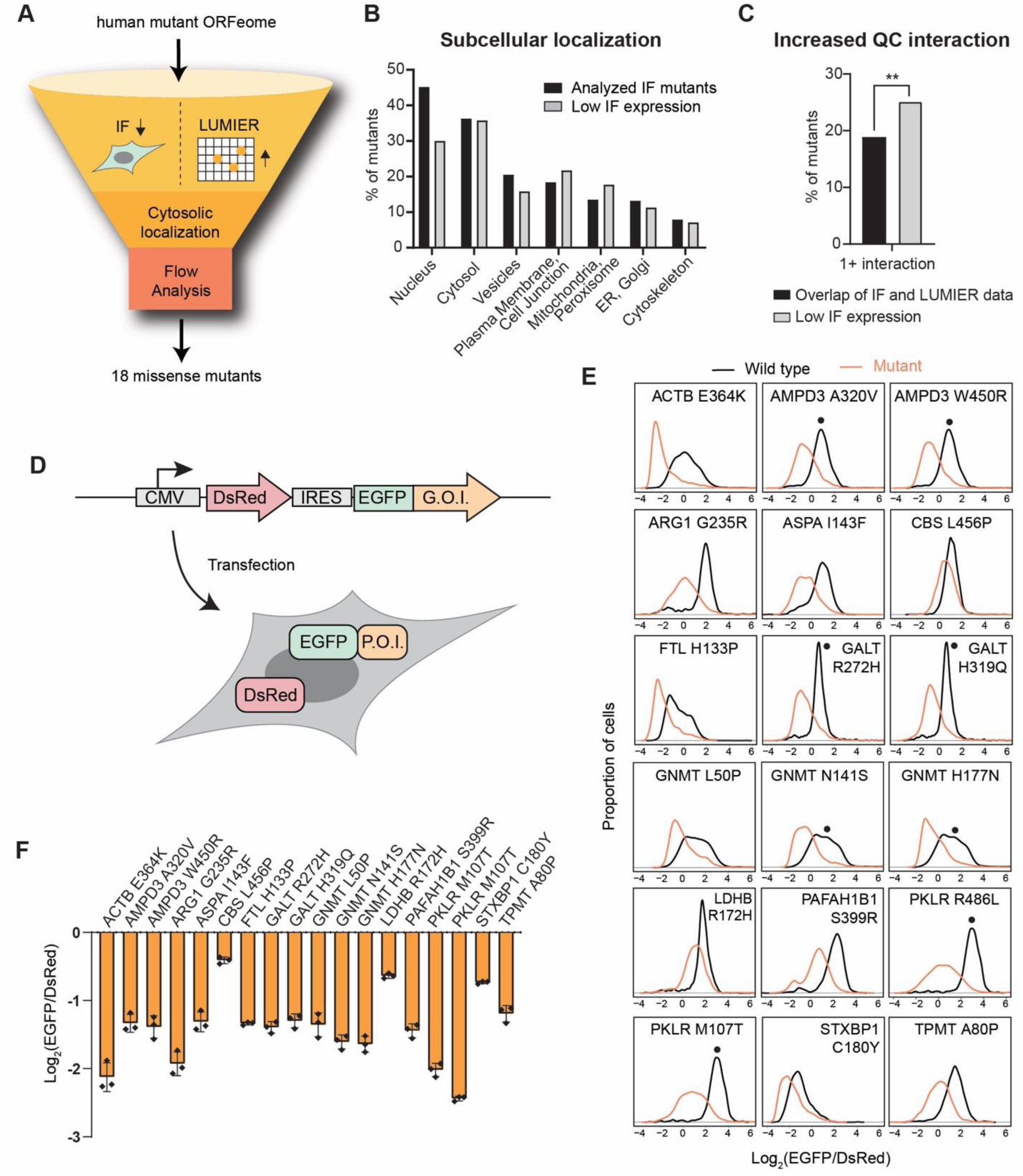
Identification of destabilizing mutations of cytosolic proteins. A) Schematic of the selection of unstable cytosolic candidate reporters with missense mutations. B) Subcellular localization of mutants assessed in a recent study and that were expressed <0.5x relative to the WT (Lacoste et al. 2023). C) Proportions of mutants assessed in the IF screen and within the LUMIER assay data (Sahni et al. 2015) that identified mutants that show increased interaction with quality control factors (QCFs). Results of Fisher’s exact test is shown (**: p-value ≤0.01). D) Schematic of the bicistronic IRES construct used for transient transfection experiments. E) Distributions of the indicated transfected reporters for the log_2_(EGFP/DsRed) signal. For genes with multiple mutants, the same WT sample was used for the comparison and are denoted with an asterisk. F) Misfolded cytosolic reporters cloned into the bicistronic IRES construct and tested for protein stability using flow cytometry. The quantitation was calculated by taking the log_2_ of the median (EGFP_MG132_/DsRed _MG132_) –log_2_ of the median (EGFP _DMSO_/DsRed _DMSO_) (n=3).

**Table 1.** Disease-associated missense mutants as model substrates to study protein quality control in the cytosol. List of cytosolic mutant substrates that were used in this study. The gene name and corresponding disease-associated protein mutation is listed. Candidates were selected based on lower levels of immunofluorescence (IF), quantitation of detection in an ELISA assay, or increased interactions with protein quality control (QC) elements (Sahni et al., 2015) all relative to the corresponding WT. Location indicates the secondary structure at the mutated residue.

| Gene | Mutation | IF | ELISA | QC | Location |
| --- | --- | --- | --- | --- | --- |
| ACTB | E364K |  |  | ✓ | B-strand |
| AMPD3 | A320V |  |  | ✓ | Helix |
| AMPD3 | W450R | ✓ |  | ✓ | Helix |
| ARG1 | G235R | ✓ | ✓ | ✓ | Helix |
| ASPA | I143F |  | ✓ |  | Helix |
| CBS | L456P |  |  | ✓ | B-strand |
| FTL | H133P | ✓ |  |  | Helix |
| GALT | R272H | ✓ |  | ✓ | Helix |
| GALT | H319Q | ✓ |  | ✓ | B-strand |
| GNMT | L50P | ✓ |  |  | Helix |
| GNMT | N141S | ✓ |  |  | Loop |
| GNMT | H177N | ✓ |  | ✓ | B-strand |
| LDHB | R172H | ✓ |  |  | Helix |
| PAFAH1B1 | S399R |  |  | ✓ | B-strand |
| PKLR | M107T |  |  | ✓ | Helix |
| PKLR | R486L | ✓ |  | ✓ | Helix |
| STXBP1 | C180Y | ✓ |  | ✓ | Helix |
| TPMT | A80P |  |  | ✓ | Helix |

### The UPS is a major pathway for the degradation of unstable variants

Since the UPS is a major degradation pathway, we first assessed whether the new model substrates are degraded in a proteasome-dependent manner. To do this, we treated HEK293T cells transfected with the mutant or WT reporter constructs with the MG132 proteasome inhibitor for 16 hours prior to flow cytometry analysis. Fluorescence levels of several WT proteins fused to GFP, such as the beta-actin (ACTB), ferritin light chain (FTL) and syntaxin-binding protein 1 (STXBP1) were markedly increased by proteasome inhibition (Figure 2A), indicating that these proteins are also degraded to an extent in our system, while all other WT proteins were not prominently affected. Importantly, levels of most mutant proteins were significantly enhanced relative to their WT counterparts upon proteasome inhibition (Figure 2A), with the exception of FTL (H133P), one of the glycine N-methyltransferase (GNMT) variants, and L-lactate dehydrogenase B chain (LDHB R172H) that was not stabilized. These results indicate that the turnover of most assessed mutants is, at least partially, dependent on degradation by the proteasome. We confirmed that a similar stabilization was observed when the irreversible proteasome inhibitor bortezomib was added to cells transfected with the different mutants (Figure 2B). Likewise, inhibition of the UBA1 ubiquitin E1-activating enzyme with TAK243 that hinders ubiquitination stabilized most assessed mutants, further suggesting that the misfolded substrates are targets of the UPS. Inhibition of neddylation with MLN4924 had a minimal impact on the stability of the misfolded reporters (Figure 2B), indicating that cullin-RING ligases that are activated by Nedd8 (Baek et al., 2021) do not play a major role in the degradation of these proteins. Several of the mutant substrates also appear to be stabilized when VCP/p97 is inhibited with CB-5083, suggesting a role for VCP in substrate remodelling or unfolding prior to degradation at the proteasome (Dai and Li, 2001; Gallagher et al., 2014; van den Boom and Meyer, 2018). In contrast, inhibition of the autophagy pathway using bafilomycin A which blocks the acidification of the lysosome did not induce a strong increase of the mutant levels (Figure 2B). Altogether, these results indicate that the UPS is the main degradation pathway that targets the assessed cytosolic mutant substrates.

**Figure 2.**
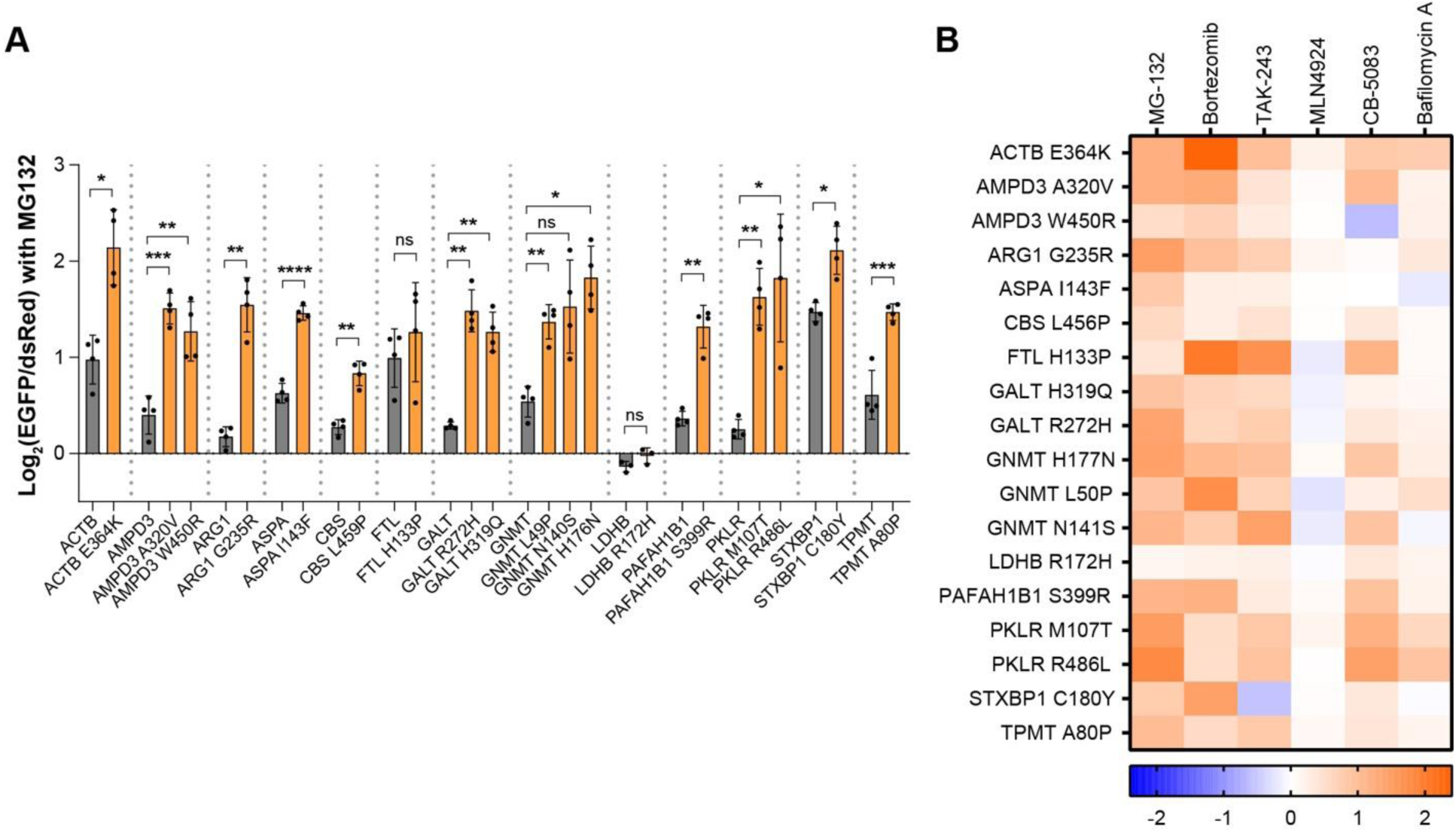
The UPS plays a major role for the degradation of cytosolic reporter with missense mutations. A) Fold change (FC) of the normalized EGFP signals of the indicated proteins upon MG132 treatment: log_2_ of the median (EGFP_MG132_/DsRed _MG132_) – log_2_ of the median (EGFP _DMSO_/DsRed _DMSO_). 32 hours after transfection, cells were treated with 5µM MG132 or DMSO for 16 hours and cells were collected 48 hours after transfection. Mean values with s.d. are shown with results of student’s t-test (n=4; p-value ns (not significant), *: p-value ≤0.05, **: p-value ≤0.01, ***: p-value ≤ 0.005, ****: p-value ≤ 0.001). B) Heatmap showing the FC of the normalized EGFP signal upon the addition of the indicated small molecule inhibitors relative to a DMSO control (a value of −2 indicates that the substrate is further destabilized and a value of 2 indicates that the substrate is stabilized).

### Unstable variants of cytosolic proteins are aggregation prone

We next used microscopy to further investigate the localization and fate of these unstable proteins in HEK293T cells. For better visualization of the reporter, we generated a stable cell line expressing an alternate dual-fluorescent reporter construct, where the mutant substrate is C-terminally fused to mClover2, and mRFP1 is used as an internal control downstream of a P2A self-cleaving peptide sequence (Figure 3A). We used one of the cytosolic model substrates, thiopurine S-methyltransferase (TPMT) and the A80P mutant, to validate the construct. Molecular dynamics simulations show that the A80P mutation is destabilizing and significantly distorts the protein structure (Rutherford and Daggett, 2008). We tested the stability of the WT and mutant in these new constructs and observed a similar marked decrease in the mutant fluorescence as with the IRES construct (Figure S1A). In addition, inhibiting proteasomal degradation in the cells stably expressing the mutant reporter resulted in significant stabilization, while not impacting the WT reporter, consistent with previous results (Figure S1B).

**Figure 3.**
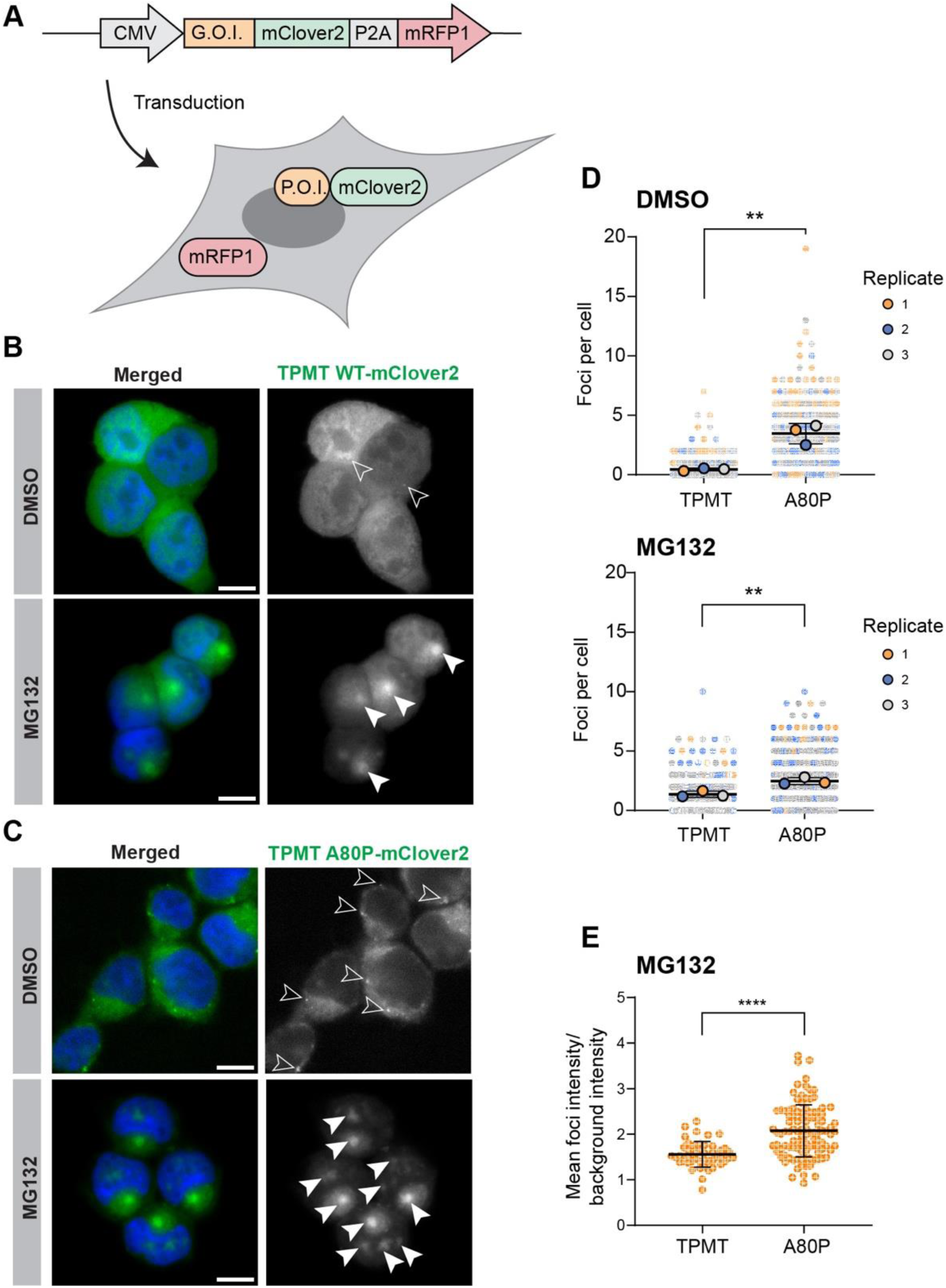
The TPMT mutant protein is recruited into foci. A) Schematic of the lentiviral bicistronic P2A construct used for microscopy experiments. B and C) Representative fluorescence microscopy images of cells transduced with TPMT-WT (B) or A80P (C) fused to mClover2-P2A-mRFP1 treated with 5 µM MG132 or DMSO for 14 hours. Hoechst-33342 was used for the staining of the nuclei. Large foci are indicated by white filled arrows and small foci with hollowed arrowheads. Scale bars: 10 μm. Note that the set up for image acquisition were identical between WT and A80P expressing cells, but brightness was adjusted for visualization, as mutant levels are lower. D) Quantitation of the number of foci per cell in three independent experiments. Individual data points are colour coded based on replicate (blue, orange, grey). The mean number of foci per cell of each replicate is denoted by a larger point outlined in black. The mean of the three replicates, S.E.M., and the results of the student’s unpaired t-test is also shown (n=3; **: p-value ≤0.01). E) Quantification of the mean foci fluorescence intensity in cells stably expressing TPMT WT or A80P-mClover2-P2A-mRFP1 upon treatment with 5 µM MG132 for 14 hours. The mean intensity of each foci per cell was averaged and compared to the background intensity in the cell. The mean across all cells with s.d. is shown and compared using a student’s unpaired t-test (n = 49 (WT), 107(mutant); ****: p-value ≤0.001).

Upon imaging, we observed that the A80P mutant forms several small cytosolic foci in untreated or DMSO-treated control cells that were not readily observed when the WT version was expressed (Figure 3B, C, S1C). Quantitation confirmed there were significantly more foci per cell on average in the A80P expressing cells (Figure 3D, S1C). When proteasomal degradation was blocked, we observed significantly larger foci in A80P expressing cells (Figure 3C, S1D). In most cells, a prominent perinuclear focus could be observed, while additional and less intense foci often appeared in the nucleus. Foci were also observed upon expression of the WT proteins (Figure 3B), but there were more foci in the A80P cells compared to the WT, and these foci were larger on average (Figure 3D, S1D). In addition, only a subset of the WT protein appeared to be recruited to the foci, while most of mutant protein was localized in foci. Indeed, the relative fluorescence intensity of the foci as compared to the cellular background signal was higher in A80P cells (Figure 3E), which suggests that, when proteasomal degradation is hindered, more of the mutant protein is being recruited into these inclusions. One possibility is that under stress conditions, such as proteasomal inhibition, the WT TPMT that is otherwise stable becomes misfolded. In contrast, the expressed mutant TPMT-A80P protein likely cannot maintain its native state in unstressed conditions and is readily recruited to large foci when proteasomal degradation is impaired.

We similarly expressed the other mutant proteins and observed the presence of large foci with all assessed protein mutants upon proteasome inhibition (Figure S1E). Foci were present in 44% to 75% of the cells (Figure S1E). Similarly, foci were also observed when the ubiquitin-activating enzyme (E1) was inhibited using TAK243 (Figure S1E, S2). The localization of these foci shows slight variations between reporters, with many showing a majority of cytosolic or perinuclear localization, while others appeared to localize to the nucleus (Figure S2). This suggests that undegraded mutant proteins may be sequestered into foci by the cells in an attempt to mitigate the aberrant effects, similar to other misfolded proteins (Johnston and Samant, 2021). Together, our results indicate that we have identified a series of cytosolic proteins that are unstable and degraded by the UPS due to missense mutations that likely induce misfolding when expressed in HEK293T cells. These new reporters can be used to study degradation QC pathways that target cytosolic proteins.

### The mutated protein TPMT A80P displays a distinct interactome

To probe for specific interactors of misfolded cytosolic proteins, we next employed the BioID proximity labelling approach (Roux et al., 2012). This method relies on an abortive biotin ligase (BirA*) fused to the protein of interest or “bait”, which leads to the biotinylation of proteins that are proximal to the bait (Figure 4A). These biotin-labelled protein interactors, or “prey”, are then pulled down with a streptavidin resin and identified using mass spectrometry. One potential advantage of this approach is that it can capture low-affinity interactions that are often lost during co-immunoprecipitation. The TPMT WT and A80P proteins were C-terminally tagged with BirA*-FLAG and the resulting fusion was stably integrated into 293 Flp-In T-Rex cells. BirA*-FLAG alone was expressed in the same cell line as a control. As expected, the TPMT A80P-BirA*-FLAG variant was expressed at lower levels than the WT protein and effected an overall lower level of biotinylation, consistent with the decreased stability of the mutant (Figure 4B). As the mutant protein is rapidly degraded by the proteasome, we also included an additional MG132 treatment group. 14 and 22 prey proteins displayed increased interactions with the mutant protein (>2-fold peptide counts) in DMSO and MG132-treated cells, respectively (Figure 4C). Many of the identified mutant-enriched preys are associated with the proteostasis network suggesting that the presence of a single missense mutation can have a profound impact both on the folding of TPMT and its interactome.

**Figure 4.**
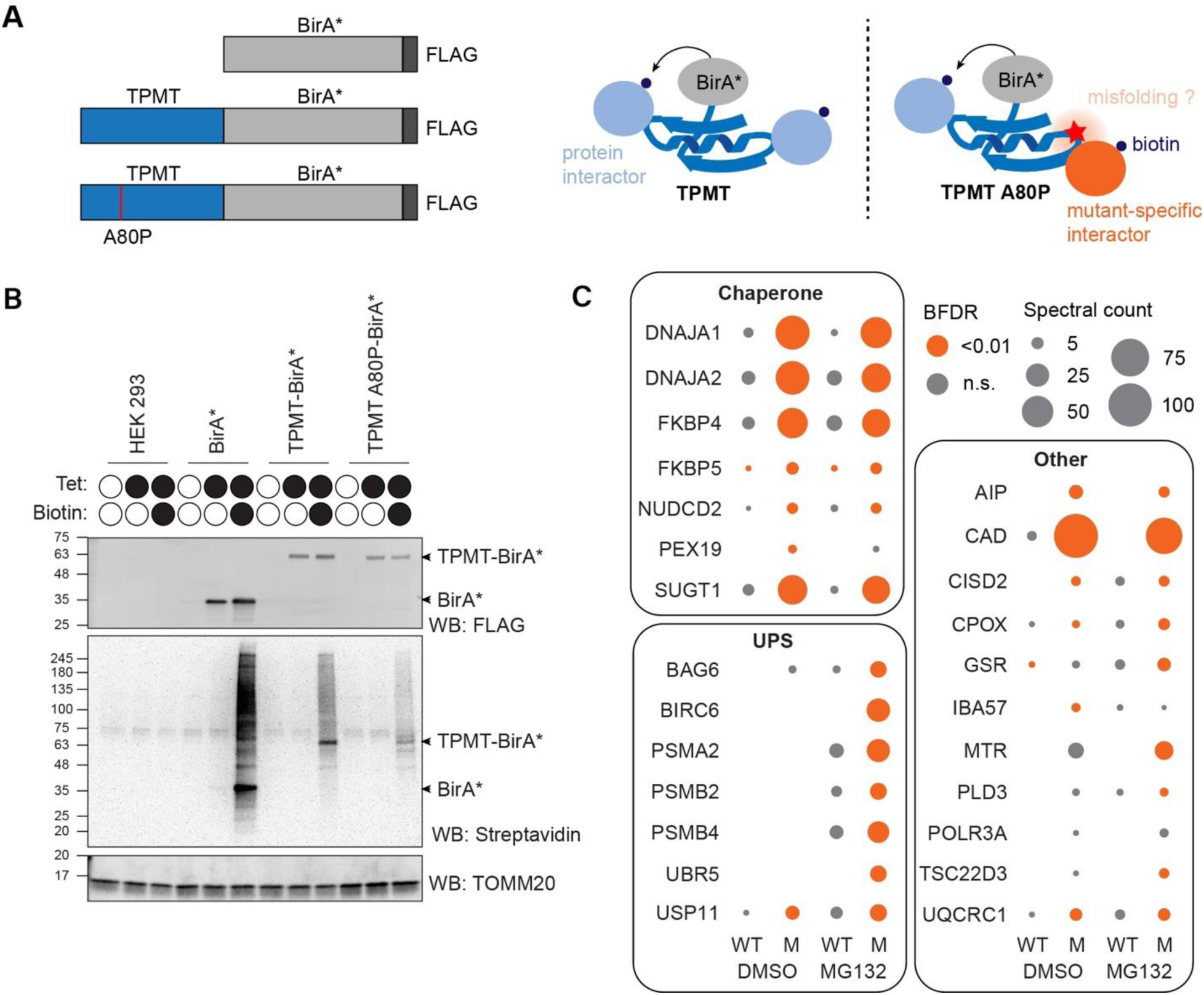
BioID proximity-labelling reveals a shift of the TPMT mutant interactome. A) Schematic of the BioID constructs (left) and representation of BioID tagging of TPMT and the misfolded variant, A80P (right). B) Western blots of the cells stably expressing the indicated baits that were treated as specified with 1 μg/ml tetracycline (Tet) for 24 h prior to addition of 50 μM biotin for 24 h. C) Dot plots representing the average numbers of spectra identified for the indicated prey proteins with the specified baits and conditions after subtracting the spectra number in the control. WT = TPMT WT-BirA*-FLAG; M = TPMT A80P-BirA*-FLAG.

Notably enriched binding to the TPMT A80P variant was observed for DNAJA1 and DNAJA2, HSP40/J-domain co-chaperones that facilitate client binding to HSP70/HSC70 (Figure 4C) (Kampinga and Craig, 2010). SGT1 and FKBP4 also displayed enrichment in the mutant BioID. SGT1 is a conserved HSP90 co-chaperone linked to degradation pathways via its interaction with SKP1 (Bansal et al., 2004; Eisele et al., 2021; Kitagawa et al., 1999). FKBP4 and FKBP5 (identified to a lower extent) are also HSP90 co-chaperones thought to act primarily as protein scaffolds (Rein, 2020; Schopf et al., 2017).

Interaction of the mutant protein with UPS proteins was even more pronounced in MG132-treated cells and included the E3 ligases UBR5 and BIRC6. UBR5 is an E3 ligase recently implicated in the quality control of newly synthesized and aggregation-prone proteins through the generation of heterotypic polyubiquitin chains that mark substrates for degradation (Yau et al., 2017). BIRC6 is an inhibitor of apoptosis and regulates cell death pathways with an additional function of acting as an E2/E3 ligase (Bartke et al., 2004). The co-chaperone BAG6 was also identified in the MG132-treated sample. BAG6 has been implicated in the quality control of proteins mislocalized to the cytosol, in the retrotranslocation of ERAD substrates, and in the selective degradation of aberrant nascent polypeptides (Hessa et al., 2011; Rodrigo-Brenni et al., 2014; Wang et al., 2011). Components of the 20S core particle (PSMA2, PSMB2, and PSMB4) were also enriched, suggesting that the mutant protein is shuttled to foci reminiscent of aggresomes (Johnston et al., 1998; Wilde et al., 2011).

### DNAJA2 stabilizes the mutated TPMT A80P reporter

To determine which identified prey protein has a dominant role in the turnover of the assessed misfolded protein, we devised a boutique genetic screen in HEK293T cells expressing the transduced TPMT-A80P-mClover2-P2A-mRFP1 reporter (Figure 5A). For all non-essential candidate hit genes associated with the proteostasis network, we designed two single guide RNAs (sgRNAs) to target an early exon of the gene that were then transfected prior to flow cytometry to measure the green-to-red fluorescence ratio. Since the plasmid containing the sgRNA of interest also contains a puromycin cassette, we could then select cells that were successfully transfected. As proof of concept, we tested the efficacy with the sgRNAs targeting DNAJA2. We saw with Western blot that one of the sgRNAs (#1) successfully targeted the DNAJA2 (Figure 5B), however, the other sgRNA (#2) was ineffective and excluded from further study. After screening the eleven hits, we saw a marked reduction of the TPMT A80P averaged signal when the co-chaperones DNAJA1 and DNAJA2 were knocked down, and an increased signal when BIRC6 was targeted (Figure 5C). We confirmed that the reduction of the reporter levels was significant (Figure 5D, S3). The knock down of additional hit genes also displayed significant changes, but below 25%. These results suggest that BIRC6 may be involved, at least partially, in the targeting of the mutated variant and that DNAJA1 and DNAJA2 are required to help slow or prevent the degradation of the mutant protein reporter.

**Figure 5.**
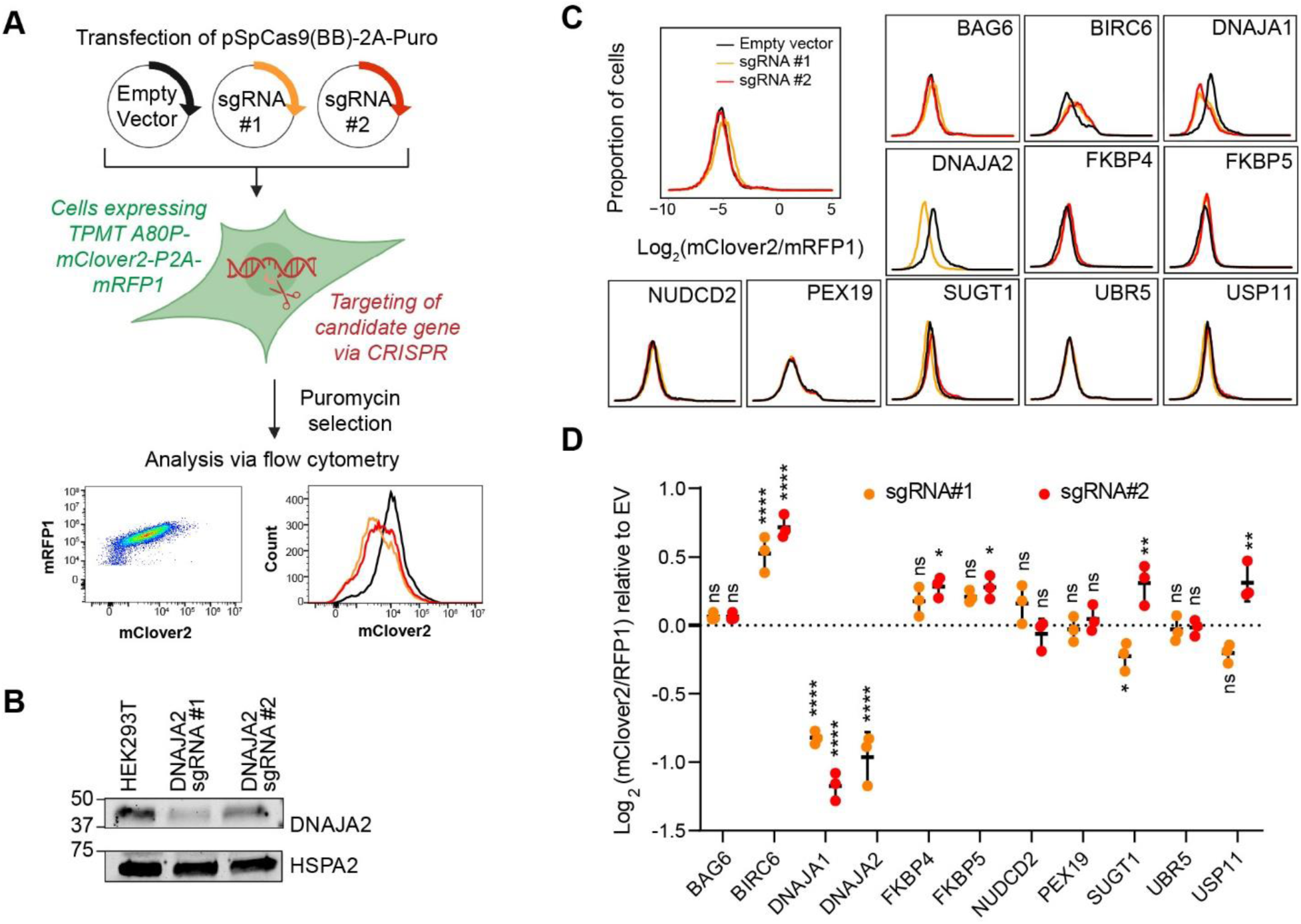
Validation of top hits from BioID reveal J-proteins as stabilizing the cytosolic misfolded substrate. A) Workflow for transfection and selection of reporter cell lines with plasmids encoding Cas9, guide RNA and a puromycin selection cassette. Assays are performed with two different sgRNAs (orange, red) and an empty vector plasmid (black) used as a control. Created with Biorender.com B) Western blots of DNAJA2 following the indicated pooled KO with the two sgRNAs. C) Representative histograms of results from screening CRISPR pooled KO of potential modulators of cytosolic misfolded proteins. D) Log_2_ of the median TPMT A80P mClover/RFP CRISPR pooled KO for each target gene relative to the empty vector negative control (log_2_ of the median (mClover_TPMT A80P_/RFP_TPMT A80P_) –log_2_ of the median(mClover _EV_/RFP_EV_)). The mean values and s.d. are shown with the results of the one-way ANOVA test, followed by the Dunnett’s multiple comparisons test against the empty vector construct (shown in details in Figure S3) (n=3; p-value ns: not significant, *: p-value ≤0.05, **: p-value ≤0.01, ***: p-value ≤ 0.005, ****: p-value ≤ 0.001).

We generated *DNAJA2* knockout (KO) clones (Figure S4A) that were then transduced with the reporters via lentivirus for flow analysis. Consistent with our previous results, the TPMT A80P reporter was further destabilized in *DNAJA2* KO cells in two independent clones (Figure 6A). Importantly, this effect was not observed with the WT TPMT protein indicating that DNAJA2 is only required to prevent degradation of the mutant protein but it is dispensable for the WT protein stability in the assessed cells. We next integrated DNAJA2-FLAG in one of the KO clones that expresses the mutant reporter and observed similar TPMT A80P levels as in *DNAJA2* WT cells when tetracycline was added to induce expression of the co-chaperone (Figure 6B and S4B). Finally, we verified that DNAJA2 displays an increased interaction with TPMT A80P compared to TPMT WT using a reverse BioID approach (Figure 6C). More specifically, the TPMT-FLAG proteins were expressed in cells stably expressing DNAJA2 fused to TurboID which allows for a shorter labelling period (Branon et al., 2018). As a negative control, we used the *Renilla* luciferase (RLuc) in parallel. The mutant protein, TPMT A80P, was found to be enriched in the DNAJA2-TurboID cells relative to the TPMT WT after the pulldown with anti-biotin. We repeated a similar experiment to confirm these results (Figure S4C). This strongly suggests that there is an increase in interaction between DNAJA2 and the mutant protein, which recapitulates what was observed in the original BioID experiment. These results show that DNAJA2 specifically interacts with the mutated protein and its presence is required to prevent a more robust degradation of the TPMT A80P mutant protein, while it is not essential for the stability of the WT protein in the assessed cells.

**Figure 6.**
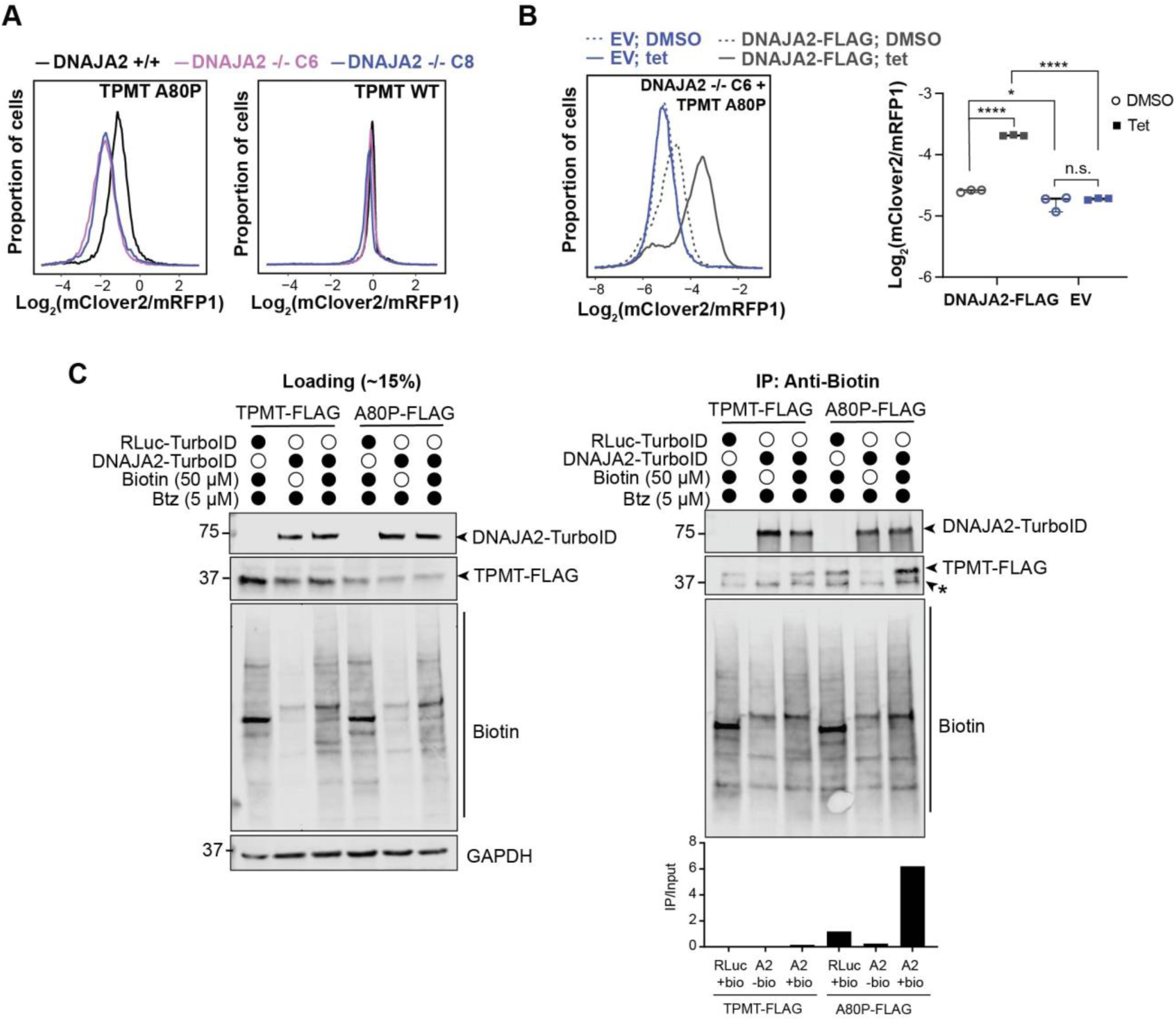
DNAJA2 stabilizes the mutant TPMT variant A80P. A) Distribution of the normalized levels of the TPMT WT and A80P reporters in two independent *DNAJA2* KO cell lines (purple and blue) and parental 293 Flp-In T-Rex cells (black). Cells were transduced with the P2A reporters. B) Distribution of the normalized levels of TPMT A80P (transduced P2A reporter) in *DNAJA2* KO cells (Flp-In HEK293, clone #6) integrated with an empty vector (EV) control or DNAJA2-FLAG. Expression of DNAJA2-FLAG was induced through the addition of tetracycline (tet) for 24 hours. The mean values and s.d. are shown with the results of the ANOVA test (n=3; *: p-value ≤0.05, ****: p-value ≤ 0.001, p-value ns: not significant). (C) Co-immunoprecipitation of biotinylated proteins in DNAJA2-BioID cells transfected with TPMT WT-FLAG and TPMT A80P-FLAG (2x the amount of DNA). Biotinylated proteins were captured using Protein G beads bound to anti-biotin antibodies. Quantitation of TPMT-FLAG signal in the IP blot relative to the lysate with the background subtracted is also shown.

### DNAJA2 has distinct clients in the cytosol

We next wanted to determine the extent of DNAJA2’s role in protein stabilization and specificity. DNAJA2 was previously shown to help target mutant CFTR for degradation at the plasma membrane, with the CHIP E3 ligase (Kim Chiaw et al., 2019). A recent study also showed that DNAJA2 can stabilize mutant p53, which normally resides in the nucleus, and slow its degradation via sequestration of the mutant (Zoltsman et al., 2023). Importantly, the role of DNAJA2 in the degradation of misfolded cytosolic proteins more broadly has not been fully investigated. Therefore, we assessed our initial panel of cytosolic missense substrates in cells lacking the co-chaperone.

We transfected the *DNAJA2* KO cells with 16 other misfolded reporters and their corresponding WT proteins for flow cytometry analysis. Interestingly, we observed that similar to our findings with TPMT A80P, two mutants were markedly destabilized in the *DNAJA2* KO: platelet-activating factor acetylhydrolase 1b regulatory subunit 1 (PAFAH1B1) S399R and pyruvate kinase (PKLR) R486L (7A; highlighted in red). In both cases, the WT PAFAH1B1 and PKLR are also mildly but significantly altered in the KO cells. Nonetheless, PAFAH1B1 S399R and PKLR R486L showed a greater decrease relative to the WT protein. In contrast, while the other PKLR variant M107T does show a significant change upon DNAJA2 KO, the change in fluorescent levels relative to the PKLR WT is very marginal (Figure 7A; highlighted in grey). This implies that certain types of mutations may induce the exposure of sites that are preferentially recognized by DNAJA2 and that this effect is not specific to the WT protein. Similarly, levels of four additional mutant reporters were not affected in the *DNAJA2* KO cells: ACTB E364K, ARG1 G235R, FTL H133P, and GNMT L50P (Figure 7A, S4; highlighted in grey). These mutations may ablate the ability of DNAJA2 to bind, as it apparent that DNAJA2 is required for the corresponding WT protein. Additionally, a large portion (9/17) of both the mutants and corresponding WT proteins were found to be present at much lower levels when expressed in *DNAJA2* KO cells (Figure 7A, S5A; highlighted in blue). In these latter cases, the presence of DNAJA2 is seemingly required for proper expression of the native proteins, whether or not these proteins carry missense mutations. Taken together, our results suggest that in some cases, DNAJA2 only “engages” some proteins when mutated to ultimately slow down the degradation of these substrates. By doing so, the co-chaperone plays a role in “buffering” proteins with mutations from the degradation machinery in the cell.

**Figure 7.**
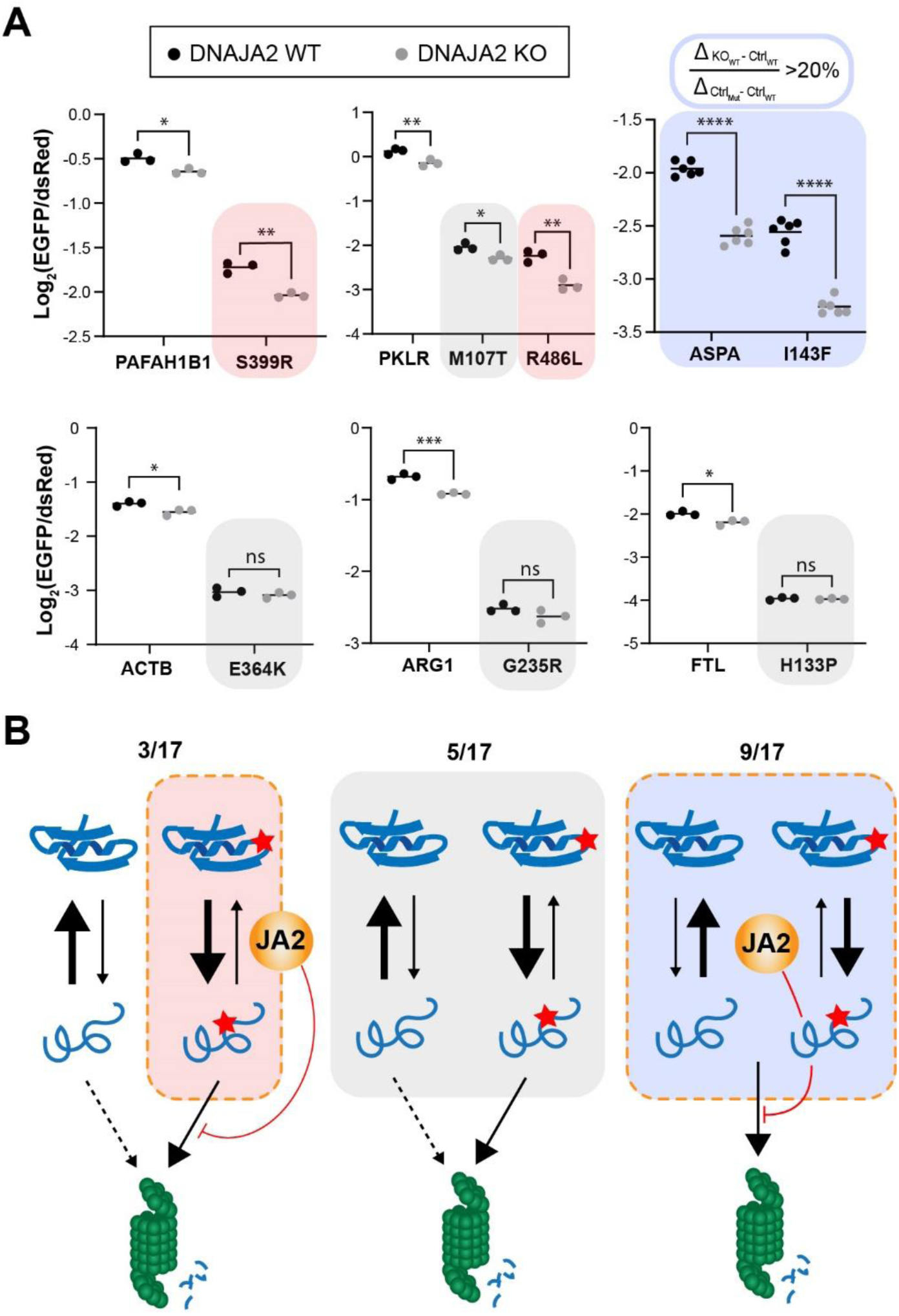
DNAJA2 is important for stabilizing WT and mutant proteins to varied extents. A) Log_2_ of the normalized EGFP signal (fold change) of transfected IRES reporters in *DNAJA2* KO cells (clone #6) or *DNAJA2* cells (293 Flp-In T-Rex). For each reporter (WT or mutant), the median of the EGFP signal normalized by the DsRed signal is log_2_ transformed. Mean values with s.d. are shown with results of student’s t-test (n=3 or n=6; *: p-value ≤0.05, **: p-value ≤0.01, ***: p-value ≤ 0.005, ****: p-value ≤ 0.001, p-value ns: not significant). To determine the cut-off for WT reporters significantly affected by DNAJA2: The fold change for each WT reporter in the DNAJA2 KO relative to the DNAJA2 control cells (Δ_KO WT – Ctrl WT_) was compared to the fold change between WT and mutant reporter in the DNAJA2 control cells (Δ_Ctrl Mut– Ctrl WT_). If Δ_KO WT – Ctrl WT_ / Δ_Ctrl Mut– Ctrl WT_ > 20%, the WT reporters were considered as significantly affected by the *DNAJA2* KO. B) Schematic of the model for the proposed action of DNAJA2 for WT and mutant proteins. DNAJA2 can buffer proteins with mutations while having a minimal effect on the WT (left panel), have no effect (middle panel), or be important in stabilizing both mutant and corresponding WT proteins (right).

## Discussion

In this study, we investigated the role of different elements of the protein quality control network that triage cytosolic missense mutant proteins. Leveraging a library of disease-associated mutants in combination with an established assay to monitor protein stability by flow cytometry (Sahni et al., 2015; Yen et al., 2008), we first identified a panel of 18 new model substrates to assess cytosolic protein quality control. We showed that the UPS plays a major role in the degradation of all of these substrates but one. To reveal how a single missense mutation on cytosolic proteins can lead to gained protein-protein interactions with the proteostasis network, we employed the BioID proximity labelling approach on the TPMT protein and its A80P variant. Chaperones and components of the UPS were found to be significantly enriched with the TPMT A80P variant in comparison to the WT protein. We then screened top interacting candidates to show that the absence of DNAJA1 and DNAJA2 increases the turnover of the TPMT A80P variant. We then further validated the involvement of DNAJA2 showing that this co-chaperone plays a role in stabilizing a subset of cytosolic proteins with missense mutations while they are dispensable in the folding of the WT proteins.

A clear observation from our study is the predominant role of the UPS in managing misfolded cytosolic proteins. The UPS remains a critical pathway in the degradation of aberrant polypeptides, offering the cell a means to control and recycle misfolded or damaged proteins. Notably, upon inhibition of the proteasome, all the assessed reporters formed foci in the cell reminiscent of aggresomes. Interestingly, inhibition of ubiquitination with TAK-243 led to the formation of similar foci. This is intriguing because HDAC6 mediates the formation of aggresome-like bodies by binding to non-degraded conjugated proteins via its ubiquitin-binding zinc finger domain (Kawaguchi et al., 2003). One possibility is that in the absence of substrate ubiquitination, neddylation may instead promote the HDAC6-dependent coalescence of the misfolded proteins or the BAG3-mediated sequestration mechanism may not require ubiquitination (Gamerdinger et al., 2011; Kim et al., 2021).

While the UPS is likely the main pathway in the degradation of the mutant substrates, proteasomal inhibition did not re-establish WT protein levels. Either because more time is required to achieve steady-state levels or because these mutant substrates are targeted for degradation via other pathways. For instance, there is cross-talk between the UPS and autophagy that has been reported for several mutant proteins (Ji and Kwon, 2017). In particular, both of these pathways play critical roles in targeting substrates associated with proteinopathies like Parkinson’s and Huntington’s diseases (Bates et al., 2015; Poewe et al., 2017). However, based on our data with Bafilomycin A, autophagy does not seem to have a dominant effect on the turnover of these cytosolic substrates. Recent studies highlight the role of mitochondria in the degradation of protein aggregates in both yeast and mammalian cells, including proteins sequestered into protein condensates (Liu et al., 2023; Ruan et al., 2017). Intriguingly, DNAJA2 along with FUNDC1 were recently shown to promote, via mitochondrial import, the degradation of TDP-43, which is associated with amyotrophic lateral sclerosis (Ma et al., 2023). The contribution of mitochondria in the degradation of misfolded cytosolic proteins would need to be more carefully evaluated in the future.

The BioID experiment revealed that numerous components of the protein quality control network are more enriched in the presence of the TPMT A80P variant. This confirms that the single point mutation induces protein misfolding. The marked enrichment of chaperones and co-chaperones with the mutant variant was independent of proteasomal degradation, whereas the biotinylation of proteins involved in the UPS was only more prominent upon MG132 treatment.

The number of spectra suggests that the interaction between the mutant protein and the molecular chaperones is more prevalent. When we assessed the potential importance of these interactions by CRISPR/Cas9 targeting, the pooled KO of *DNAJA1* and *DNAJA2* led to lower levels of the TPMT A80P variant, while other targeted chaperone genes did not impact the reporter degradation. We cannot exclude the role of these other molecular chaperones, as they may also have overlapping and redundant functions in the cell with negligible impact on substrate turnover when singly targeted. This may also explain why the pooled KO of *DNAJA1* or *DNAJA2* only has a minor impact on the reporter turnover.

To gauge the impact of DNAJA2 on cytosolic protein quality control, we assessed our panel of model substrates in the knock out cells. In many cases, the absence of the co-chaperone in the KO cells led to markedly lower levels of both the WT and mutant proteins. We posit that DNAJA2 is required, in these conditions, for the folding of these proteins to their native states and to avoid their targeting to the 26S proteasome (Figure 7B, right panel). The DNAJA2-dependence may be exacerbated by the fact these reporters are overexpressed upon transient transfection. In several other cases, the absence of DNAJA2 had no or little effect, suggesting it was dispensable for the folding of these proteins, and the co-chaperone did not bind to these mutated variants (Figure 7B, middle panel). Finally, in the last cases, DNAJA2 provides a “buffering” capacity for proteins with mutations, which leads to increased stability of the mutant protein with little impact on the WT proteins (Figure 7B, left panel). Hsp90, another major chaperone family, has a well-established role in buffering genetic variation, both in model organisms and in the context of human diseases (Queitsch et al., 2002; Rutherford and Lindquist, 1998). Hsp90 can stabilize proteins with altered conformations that can arise from mutations, resulting in reducing the severity of the observed phenotype (Karras et al., 2017). This stabilization often involves the folding and signalling of proteins, such as kinases and hormone receptors, which are susceptible to conformational changes and mutations. Increasing the copy number of the gene that encodes for HSP70 in *D. melanogaster* has also been shown to reduce the effect of stress-induced developmental defects (Roberts and Feder, 1999). It is possible that the HSP70 machinery, in combination with DNAJA2, could similarly buffer genetic alterations by providing a first line of defense against misfolded proteins. The buffering capacity of this chaperone could serve as a temporary holding mechanism providing the cell with time to refold or process the mutant proteins, thereby reducing the immediate impact of mutations on cellular phenotypes.

Our findings underscore the pivotal role of the UPS in managing misfolded cytosolic proteins while highlighting the importance of DNAJA2 in stabilizing cytosolic proteins in their native and aberrant forms.

## Acknowledgements

We are grateful to Andrew Johnson and UBCFlow core for their help with flow cytometry experiments and members of the Mayor lab for discussions, especially Katharina Schatz for preliminary work for this study.

## Authors’ Contributions

H.A.B., J.P.B., V.C., B.R., M.T. and T.M. designed the experiments. H.A.B., J.P.B., V.C, A.M., S.K., and H.E. performed and interpreted the experiments. F.A.Y carried out the mass spectrometry analysis. J.L. carried out the original IF analysis of mutants. G.C. and S.C. contributed to the reagents. T.M., B.R., and M.T. supervised the research helped interpret the results. H.A.B. and T.M. wrote the manuscript in consultation with the co-authors.

## Methods

### Selection of mutants

Immunofluorescence (IF) images of HeLa cells transfected with C-terminally FLAG tagged proteins in HeLA cells (Lacoste et al., 2023) were first assessed manually to identify 369 mutant proteins not expressed. We then reanalyzed images via a python script to look at the signal intensity of mutant relative to WT, and identified 659 mutants that had <0.5 expression relative to WT were selected. Additional candidates were selected based on enriched interactions with quality control factors and reduced levels in an ELISA assay (Sahni et al., 2015). Mutants that are cytosolic were chosen for further analysis. Candidates with incorrect sequences were removed from the pool of mutants. For cell localization, Human Protein Atlas data was used. A few selected mutants were not properly characterized in the Human protein Atlas, but had GO terms associated to cytosolic localization; their cytosolic localization was verified by microscopy (Figure S2). All proteins were sequenced to ensure they were correctly annotated.

### Plasmids used in this study

All plasmids generated in this study are listed in Table S2. All WT and mutant variants from the hmORFeome1.1 were subcloned into the MSCV-CMV-DsRed-IRES-EGFP-DEST (Yen et al., 2008) or the pLenti6.2-ORF-mClover2-P2A-mRFP1 destination vector using the LR II Clonase Mix (Invitrogen) and following manufacturer’s instructions.

The pDEST-N-BirA* plasmid used in the BioID experiment. TPMT and TPMT A80P were subcloned into this vector using the the LR II Clonase Mix (Invitrogen) and following manufacturer’s instructions.

To generate the plasmids used for KO of top hits from the BioID results, first sgRNAs were selected using CHOPCHOP v. 3 (Table S2) and were ordered from IDT with BbsI flanking sites for cloning into either pSpCas9(BB)-2A-GFP (PX458) (Addgene #48138 for single cell sorting or pSpCas9(BB)-2A-Puro (PX459) V2.0 (Addgene #62988) for puromycin selection (Ran et al., 2013). All constructs were confirmed by sequencing.

The pCDNA5-FRT/TO-FLAG-DNAJA2 plasmid used in the rescue experiment was generated by amplifying the ORF of DNAJA2 with flanking XhoI/BamHI restriction sites and ligating into pGC-DNA5 (Kozak) FRT/TO Flag-HA (Table S2).

### Cell tissue culture, transfection and transduction

HEK293T cells were cultured in Dulbecco’s Modified Eagle Medium (DMEM), high glucose (Gibco) supplemented with 10% fetal bovine serum (FBS) (Gibco), 1x penicillin-streptomycin (Gibco) at 37°C and 5% CO_2_. For initial experiments, the proteasome inhibitors MG132 and Bortezomib were used at a final concentration of 5 μM for 16 hrs. 2.5 μM TAK243, 1 µM MLN-4924 and 1 nM bafilomycin A1 were added for 8 hours. The Hsp90 inhibitor Geldanamycin and the Hsp70 inhibitor Ver-15508 were used at a final concentration of 1.5 μM for 16 hours and 2.5 nM for 8 hours. FuGENE 6 (Promega) was used for all transfections according to the supplied protocol. To produce retroviruses 7×10^5^ HEK293T cells were seeded in 6 cm plates and transfected with psPAX2 (0.75 µg), pMD2.G (0.25 µg) and pLenti6.2-ORF-mClover2-P2A-mRFP (1 µg). Virus was removed from packaging cells 24 hours after transfection, passed through a 0.45 µm filter, and added to cells to be infected. To generate stable reporter cell lines, HEK293T cells were transduced with retroviruses carrying mutant or WT genes in the pLenti6.2-ORF-mClover2-P2A-mRFP plasmid supplemented with 4 μg/ml of polybrene (EMD-Millipore). Cells were selected with 6 μg/ml of blasticidin (Invitrogen) and mixed transduced cell population were used. For the BioID, 293 T-Rex Flp-In cells were co-transfected with pOG44 (Flp recombinase expression vector) and a pcDNA5-based FRT/TO BirA*-FLAG expression vector, containing the coding sequence for the BirA* protein alone, the TPMT WT-, or the TPMT A80P-tagged fusion protein. Following transfection, cells were selected using 5 μM hygromycin until colonies on the plate had expanded. The dish was split in half, into “population A” and “population B”. For pooled KO experiments, HEK293T TPMT A80P-mClover2-P2A-mRFP1 cells were transfected with the PX459 constructs and selected with 8 μg/ml puromycin for 2 days. For KO generation, 293 T-Rex Flp-In cells were transfected with the PX458 construct, harvested using trypsin after 48 hours and resuspended in 1% FBS in 1XPBS. GFP-positive cells were sorted into a 96 well plate which contained 50% fresh DMEM, 20% FBS, and 30% used pre-conditioned medium (collected from plates growing cells at 50% confluency and filtered through a 0.45µm filter). Once single colonies were identified, cells were expanded and clones were verified via Western blots. For the rescue experiments, 293 T-Rex Flp-In *DNAJA2* KO #6 was transduced with pLenti6.2-TPMT A80P-mClover2-P2A-mRFP1 and selected using 5 μM hygromycin. Cells were then co-transfected with pOG44 and the pCDNA5 FRT/TO FLAG-DNAJA2 plasmid or pCDNA5 FRT/TO Empty as the control. Cells were selected with hygromycin until colonies on the plate were separated into “population A” and “population B”. Tetracycline (1 µg/ml) or DMSO was added for 24h prior to flow analysis. For the reverse TurboID experiments, the stable cell lines were generated by co-transfecting 293 T-Rex Flp-In cells with pOG44 and RLuc-TurboID-FLAG or DNAJA2-TurboID-FLAG. Cells were selected with hygromycin. Cell lines were routinely tested for mycoplasma contamination and to our knowledge, no cell lines used in this study were contaminated.

### Flow Cytometry Analysis

For all assays using the pMCV expression plasmids flow cytometry was performed 48 hours post transfection. Flow analysis for small molecule inhibitor treatments in HEK293T cells were performed in the Attune NxT Flow Cytometry System (ThermoFisher). For all assays with the stable cell lines generated using the pLenti6.2-ORF-mClover2-P2A-mRFP vector and all experiments with the DNAJA1 and DNAJA2 KOs, cells were analyzed using the Cytoflex (Beckman-Coulter). For each analysis 10,000 events were typically collected. All data was analyzed using FlowJo. To calculate the relative abundance in each experiment, the log_2_ of the median (EGFP/dsRed) or log_2_ of the median (mClover2/mRFP1) signals of individual cells was used.

### Fluorescence microscopy and quantitation

Cells stably expressing protein of interest fused to mClover2-P2A-mRFP were seeded and grown on poly-D-lysine coated coverslips in 6-well plates until 50% confluency. Cells were then treated with either MG132 (5 µM), TAK243 (2 µM), DMSO, or left untreated for 14h (n=3). Cells were washed once with 1XPBS then fixed and stained with Hoechst 33342 in 3% paraformaldehyde for 15 min. Coverslips were washed with 1x PBS and then mounted onto slides using Dako mounting medium (Agilent). Cells were imaged using a 63x oil lens. Images were analyzed using Zen Blue (Zeiss), FIJI (Schindelin et al., 2012), and CellProfiler 4.2.6 (Stirling et al., 2021). For the experiment outlined in Figure 3B-E and S1A-B, the experiment was repeated in triplicate and a minimum of 10 fields of view were selected at random per slide and used as input for analysis in CellProfiler. Number of total cells analyzed by the pipeline: TPMT Untreated = 287; A80P untreated = 244; TPMT DMSO =245; A80P DMSO = 258; TPMT MG132 = 245; A80P MG132 = 407. For Figure S1C, a minimum of 5 fields of view were selected. Nuclei were counted manually, then cells with foci that appeared to be cytosolic, nuclear, or both were also counted manually. DMSO treated cells: n=145-305. MG132 treated cells: n=87-192. TAK243 treated cells: n=59-171.

To count the total number of foci per cell, a pipeline on CellProfiler was developed to identify the focus of each cell with a pixel diameter range of 5-30. The number of foci per cell and the area of each focus were measured. The same parameters were applied across all replicates (n=3) for the MG132, DMSO, and untreated conditions. The statistics were calculated and visualized using GraphPad Prism version 10.1.2.324. To determine the signal intensity of the foci relative to the background cytosolic signal, a separate feature of the cytosol with the foci subtracted was created (“CytoNoFoci”) within the same pipeline. The average of the mean intensity of all foci per individual cell was also calculated (“MeanAllFoci”). The mean intensity of “CellNoFoci” was also calculated. To calculate the ratio between the average foci signal and the cytosolic background, “MeanAllFoci”/ “CytoNoFoci” was calculated per cell. Cells without foci were excluded from the analysis.

### Western Blots

Samples were loaded onto 4–20% Mini-PROTEAN® TGX™ Precast Protein Gels (Bio-Rad) and transferred onto a 0.45 µm nitrocellulose membrane using the TransBlot® Turbo™ Transfer System (Bio-Rad). Samples were blocked in 5% powdered milk + 1x TBS-T. To wash, 1x TBS-T was used. Blots were imaged and quantified using the Odyssey (Li-Cor). The following primary antibodies were used at the specified dilutions and were diluted in 5% powder milk in 1x TBS-T: anti-DNAJA2 (LS-C115612-100 - Lifespan Biosciences; 1:1000), anti-FLAG (PA1-984B, Invitrogen, 1:1000), anti-GAPDH (14C10 - Cell Signaling Technology; 1:2000), anti-HSPA2 (HPA000798 – Atlas Antibodies; 1:2000), anti-TOMM20 (sc-11415, Santa Cruz; 1:5000). The following secondaries were used at the specified dilutions and were diluted in 5% powder milk in 1x TBS-T: IRDye 800CW Streptavidin (Li-Cor; 1:10000) to visualize biotin, IRDye 800CW Goat anti-Mouse (Li-Cor; 1:10000), IRDye 680RD anti-rabbit (Li-Cor; 1:10000).

### BioID

Stable isogenic cell pools expressing BirA*-FLAG, TPMT-BirA*-FLAG, or the A80P-BirA*-FLAG bait proteins were expanded to 5 x 15 cm^2^ plates at ∼80% confluency. Expression of the fusion protein was induced by the addition of 1 µg/ml tetracycline and 50 µM biotin (final concentration) to the culture media for 24 hrs. Cells were harvested, pelleted and washed in cold 1XPBS. Pellets were snap frozen in LN_2_ and stored at −70℃. Cells were thawed and lysed in 1 ml of Triton lysis buffer (50mM Tris pH 7.5, 150mM NaCl, 1% Triton) with cOmplete, EDTA-free Protease Inhibitor Cocktail (Roche) on ice for 30 min. Samples were clarified by spinning (4000 rpm, 4℃, 10 min) and soluble supernatant was added to 30 µl of High Capacity Streptavidin Agarose (Pierce) pre-equilibrated in Triton lysis buffer on an end-over-end rotator at 4℃ for 3 h. Beads were transferred to spin columns (Pierce™ Spin Columns - Snap Cap #69725) and washed (2×1 ml washes in lysis buffer). Samples were washed with 0.5% SDS in 1XPBS. Following SDS wash, the remainder of protocol was done at RT. Samples were washed 2x with 1 mL of PBS SDS solution. Beads were incubated with 200 µl of 100 mM DTT in PBS SDS solution for 20 min at RT. Beads were washed 10x in 1 mL of UC buffer (6M urea, 100 mM Tris-HCl pH 8.5) then incubated with 200 µl UC buffer with 50 mM iodoacetamide for 20 min at RT in the dark. Samples were washed 10x more in 1 mL of UC buffer then 3x in 1 mL of water. Each sample was digested on beads in 100 µl of 50 mM ammonium bicarbonate with 1 µg trypsin (Promega) in spin cap tubes sitting in fresh mass-spectrometry grade Eppendorf tubes, sealed with parafilm. Peptides were collected in a fresh MS tube by centrifugation (1000g, 1 min) then 100 µl Ammonium bicarbonate was added to beads and collected again by centrifugation as above. Samples were then acidified prior to passing them through stage tips for MS analysis.

### Mass spectrometry

High performance liquid chromatography was conducted using a 2 cm pre-column (Acclaim PepMap 50 mm x 100 um inner diameter (ID)), and 50 cm analytical column (Acclaim PepMap, 500 mm x 75 µm ID; C18; 2 µm; 100 Å, Thermo Fisher Scientific, Waltham, MA), running a 120 min reversed-phase buffer gradient at 225 nl/minute on a Proxeon EASY-nLC 1000 pump in-line with a Thermo Q-Exactive HF quadrupole-Orbitrap mass spectrometer. A parent ion scan was performed using a resolving power of 60,000, then up to the twenty most intense peaks were selected for MS/MS (minimum ion count of 1,000 for activation) using higher energy collision induced dissociation fragmentation. Dynamic exclusion was activated such that MS/MS of the same *m/z* (within a range of 10 ppm; exclusion list size = 500) detected twice within 5 sec were excluded from analysis for 15 sec. For protein identification, Thermo .RAW files were converted to the .mzXML format using Proteowizard (Kessner et al., 2008), then searched using X!Tandem (Craig and Beavis, 2004) and COMET (Eng et al., 2013) against the Human RefSeq Version 45 database (containing 36,113 entries). Data were analyzed using the trans-proteomic pipeline (TPP)(Deutsch et al., 2010) via the ProHits software suite (v3.3)(Liu et al., 2010). Search parameters specified a parent ion mass tolerance of 10 ppm, and an MS/MS fragment ion tolerance of 0.4 Da, with up to two missed cleavages allowed for trypsin. Variable modifications of +16 for M and W, +32 for M and W, +42 for N-terminus, and +1 for N and Q were allowed. Proteins identified with an iProphet cut-off of 0.9 (corresponding to ≤1% FDR) and at least two unique peptides were analyzed with SAINT Express v.3.6 (Teo et al., 2014). 10 control runs (from cells expressing the FlagBirA* epitope tag alone) were collapsed to the two highest spectral counts for each prey and compared to the experimental data, consisting of two biological replicates (each analyzed with two technical replicates). High confidence interactors were defined as those with bayesian false discovery rate (BFDR) ≤0.01.

### Reverse TurboID pulldown

DNAJA2-TurboID and RLuc-TurboID cells were seeded at 3.2×10^5^ cells per 60 mm dish. Tetracycline (1 µg/ml) was added to the dish, and 24h after addition, cells were transfected with TPMT-FLAG or TPMT A80P-FLAG with FuGENE 6 (Promega). 24 h post transfection, cells were treated with additional tetracycline (1 µg/ml). 47 h post transfection, biotin (50 µM) and bortezomib (5 µM) were added for 1 h. Cells were washed with 1X PBS and were resuspended in 1 mL of 1X Triton Lysis Buffer (50mM Tris pH 7.5, 150mM NaCl, 0.5% Triton and cOmplete, EDTA-free Protease Inhibitor Cocktail (Roche)). Samples were left on ice for 15 min and lysed using a water bath sonicator with ice water for 30s on, 30s off 2x on high. Total cell lysate was collected at this point. Samples were centrifuged at 21,000 rcf at 4°C for 20 min. Anti-biotin antibody was added to the cell lysate to mix overnight at 4°C. 20 µL of the Protein G agarose bead (Millipore) slurry was washed and added to the antibody-lysate mixture and incubated for 2 h at 4°C. The flow through was collected and the beads were washed twice with 1X Triton Lysis Buffer and once with 1X PBS. To elute the biotinylated proteins, 40 µL of 2X Lammeli Buffer was added to the beads and heated for 5 min at 96°C.

## Data Availability

The raw mass spectrometry data has been uploaded to MassIVE (http://massive.ucsd.edu/) with the accession number MSV000094061. The data that support the findings of this study are also available from the corresponding author upon request.

## Supplemental Tables

**Supplemental Table 1.** Further details about the cytosolic mutant proteins that were used in this study including the gene name, protein, nucleotide mutation, protein mutation, mutation location (based on the available structural or predicted data via Uniprot), protein localization (Protein Atlas and Uniprot), and associated disease (HMGD).

**Supplemental Table 2.** List of plasmids and sgRNAs used in this study.

**Supplementary Table 3.** SAINT analysis of BioID-MS experiment.

